# simmr: An open-source tool to perform simulations in Mendelian Randomization

**DOI:** 10.1101/2023.09.11.556975

**Authors:** Noah Lorincz-Comi, Yihe Yang, Xiaofeng Zhu

## Abstract

Mendelian Randomization (MR) has become a popular tool for inferring causality of risk factors on disease. There are currently over 45 different methods available to perform MR, reflecting this extremely active research area. It would be desirable to have a standard simulation environment to objectively evaluate the existing and future methods. We present `simmr`, an open-source software for performing simulations to evaluate the performance of MR methods in a range of scenarios encountered in practice. Researchers can directly modify the `simmr` source code so that the research community may arrive at a widely accepted framework for researchers to evaluate the performance of different MR methods.

## 1 INTRODUCTION

Mendelian Randomization is a genetic instrumental variable method that has become a popular statistical tool for estimating causal effects of risk factors on disease (Sanderson et al., 2022a; Richmond and Smith, 2022). The validity of inferences made using MR relies heavily on the satisfaction of three primary assumptions (de Leeuw et al., 2022):the genetic instruments are (i) strongly associated with the exposure(s) (Davies et al., 2015; Sanderson et al., 2021), (i) not associated with the outcome conditional on the exposures (Hemani et al., 2018), and (iii) not associated with any confounders of the exposure(s)-outcome relationship(s) (Morrison et al., 2020). Violation of any of these assumptions can lead to bias in causal estimation, therefore inflating false positive and/or false negative rates (Bowden et al., 2015). Three additional challenges are present in MR analyses: (a) some individuals may have been present in both the exposure(s) and outcome GWAS which leads to sample overlap bias (Burgess et al., 2016a), (b) applying strict IV selection criteria to satisfy assumption (i) above can introduce a ‘winner’s curse’ bias (Jiang et al., 2022b), and (c) strong LD between IVs (Gkatzionis et al., 2023), and its imprecise estimation using reference panels, can drastically inflate false positive rates (Gleason et al., 2021).

At least 45 statistical methods for performing MR have been introduced in the literature to address different subsets of assumptions (i)-(iii) and scenarios (a)-(c) in either the single-exposure (univariable) or multiple-exposure (multivariable) settings (Boehm and Zhou, 2022). Clearly, the literature is saturated with a variety of MR estimators. In each of the initial studies introducing these methods, simulation was performed to evaluate the performance of new and existing methods. The variability of simulation settings reported in the literature is vast. Some use 50 or fewer SNPs to explain most of phenoptypic (Verbanck et al., 2018; Zhu et al., 2021) or genetic (Lin et al., 2023) variance while others have used hundreds (Lorincz-Comi et al., 2023) or even thousands (Mounier and Kutalik, 2023) of SNPs. Additionally, some have used GWAS sample sizes no larger than 500 (Deng et al., 2022) while others have used GWAS sample sizes no less than 100k (Xu et al., 2021). Nearly all multivariable MR simulation sets do not accurately reflect reality. For example, some have 50 or less IVs in simulation (Lin et al., 2023; Sanderson et al., 2019, 2022b), but experience with real data analyses with multiple exposures suggest that hundreds or thousands of SNPs meeting the valid IV conditions may be identified (Davies et al., 2019; Lorincz-Comi et al., 2023). Since weak IV bias generally becomes more severe as more IVs are used (Davies et al., 2015; Lorincz-Comi et al., 2023), MVMR simulations in which very few IVs are used may unrealistically reflect optimal performance of MVMR methods which may break down in practice. Some authors have additionally modelled highly phenotypically correlated phenotypes that are completely independent genetically (Lin et al., 2023; Grant and Burgess, 2021), a scenario unlikely to be encountered in practice and one that does not require MVMR methods.

Some of these issues arise from researchers reproducing the unrealiastic simulation settings of others as described above (e.g., (Zhu et al., 2021) reproduced (Verbanck et al., 2018); (Lin et al., 2023) reproduced (Grant and Burgess,2021)). Other challenges may exist because of a lack of a unified framework for performing simulations in MR. Both explanations have produced a literature of results potentially sensitive to the unique simulation conditions that may or may not mirror reality. Indeed, some independent simulations intended to mimic the same reality with nearly identical reported simulation settings have even produced conflicting results. As examples, the literature contains some conflicting evidence about the bias in dIVW in the absence of sample overlap and horizontal pleiotropy ((Mounier and Kutalik, 2023) and (Ye et al., 2021)); the power of MR-Robust in the presence of 50% invalid IVs, being either less than 20% or greater than 80% ((Lin et al., 2023) and (Slob and Burgess, 2020)); or MR-RAPS having Type I error greater than 80% or controlled at 5% in the presence of horizontal pleiotropy ((Qi and Chatterjee, 2021) and (Xue et al., 2021)).

We introduce `simmr`, an open-source and flexible tool for generating data to use in MR simulations. The last time a tool such as this was introduced was `MR_predictor` 10 years ago (Voight, 2014), however `MR_predictor` cannot accommodate horizontal pleiotropy, sample overlap, multiple exposures, weak instruments, correlated IVs, imprecise LD estimation, or winner’s curse. `simmr` can accommodate each of these and even more scenarios simultaneously in either the UVMR or MVMR settings. With no modern standard set in the literature, the purpose of the open-source `simmr` software is to begin the construction of a standard simulation environment in which fair comparisons can be made between competing MR estimators in different settings that are encountered in practice.

## 2 SIMULATION MODELS

The simulation data-generating process is based on the directed acyclic graph (DAG) in Figure 1 using the input parameters displayed in Figure 2. That is, simulation data are generated under the following models:

**FIGURE 1.**
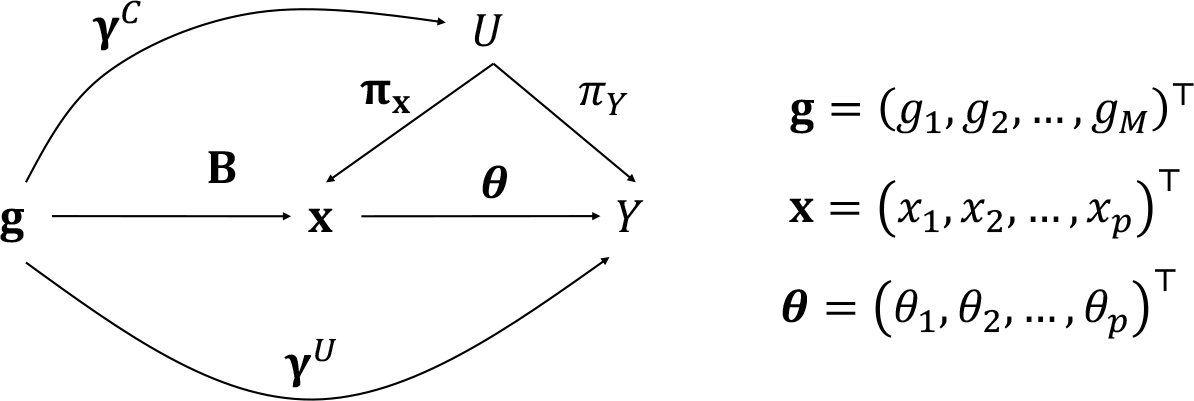
This DAG is used to produce the simulation models described in the text. Users of simmr can modify all parameters that are present in this DAG.

**FIGURE 2.**
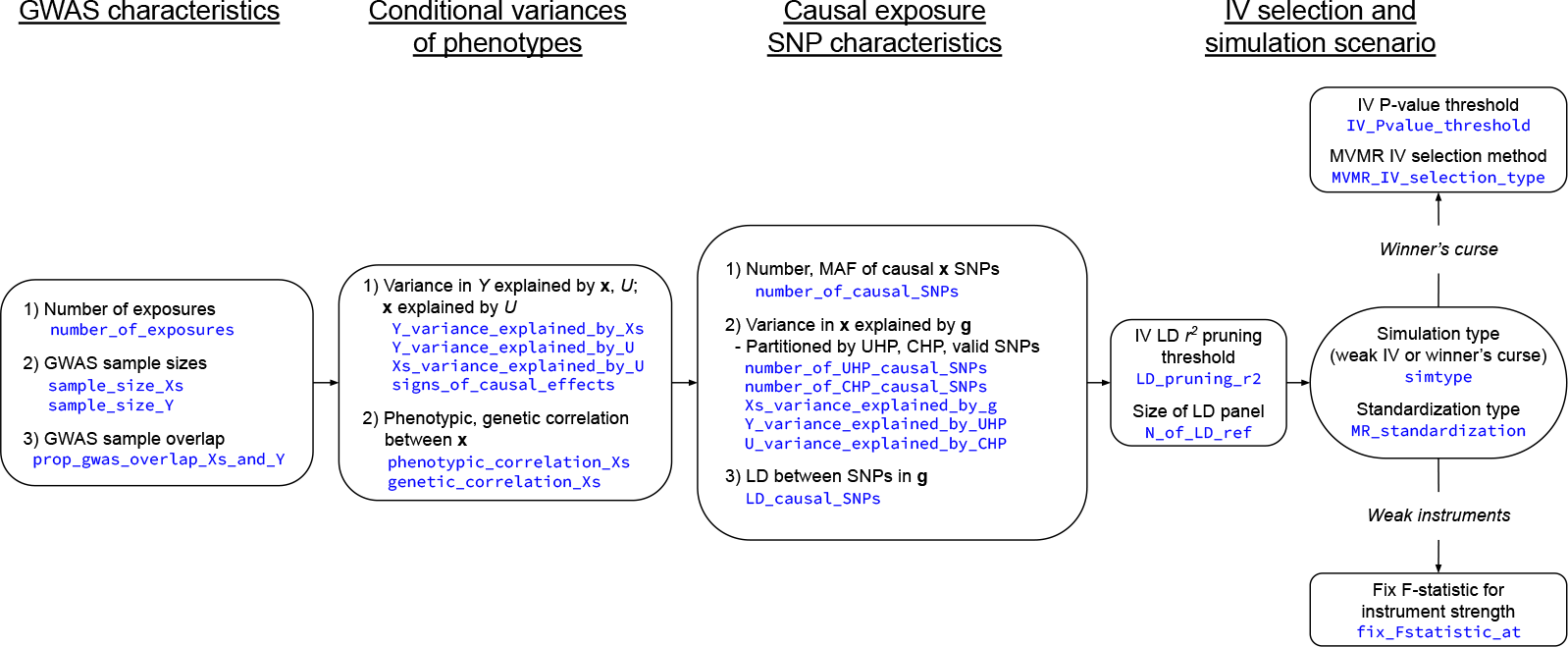
This figure provides a conceptual overview of the `simmr` data-generating process. First, users specify the sizes of exposure and outcome GWAS sample sizes and their degree of overlap. Next, users specify the true causal model by changing variances in each phenotype explained by each other and by the SNPs that are causally related to the exposures. Next, the user specifies the particular characteristics of the causal SNPs, inputting the degree of UHP, CHP, and LD between them. Finally, the user can perform IV selection to evaluate winner’s curse or weak instrument bias. `simmr` commands are presented in blue text underneath the description for each parameter. Users can view a summary of the outputted data, in the format of the right panel in Figure 3, simply by executing `plot_simdata()` after `generate_data()`.

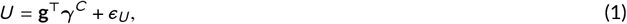

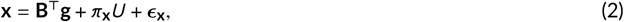

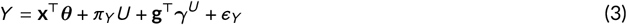

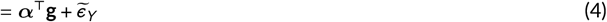

where *U* is a confounder of the relationship between exposure(s) **x** and outcome *Y*, ***θ*** represents the corresponding causal effect(s), **B** represents true associations between **g** and **x**, and ***γ***^*C*^ and ***γ***^*U*^ are respectively CHP and UHP effects. Users of `simmr` can fix the variance explained in each phenotype by the others and partition the variance in **x** explained by **g** into effects from UHP, CHP, valid, and weak SNPs using the procedure in Algorithm 1.

Figure 3 shows an example of data generated under the above models alongside computation times for different combinations of GWAS sample overlap, GWAS sample sizes, and numbers of exposure SNPs. This set of SNPs can then be reduced by the user-specified significance and LD pruning thresholds (Dudbridge and Newcombe, 2016). Users also have the option to perform IV pruning such that a specific F-statistic for instrument strength (Burgess et al., 2011) is achieved for each exposure. Some MR methods use the correlation matrix Ω among GWAS estimation errors across cohorts to correct for weak instrument bias (e.g., (Lin et al., 2023), (Lorincz-Comi et al., 2023), (Cheng et al., 2020), (Mounier and Kutalik, 2023)). These methods calculate Ω using non-significant GWAS summary statistics (Lorincz-Comi et al., 2023; Mounier and Kutalik, 2023; Zhu et al., 2015; Zhao et al., 2020b) or LD score regression (Lin et al., 2023; Cheng et al., 2020). `simmr` will directly calculate Ω using the methods in (LeBlanc et al., 2018) without any estimation error and return it to the user.

**FIGURE 3.**
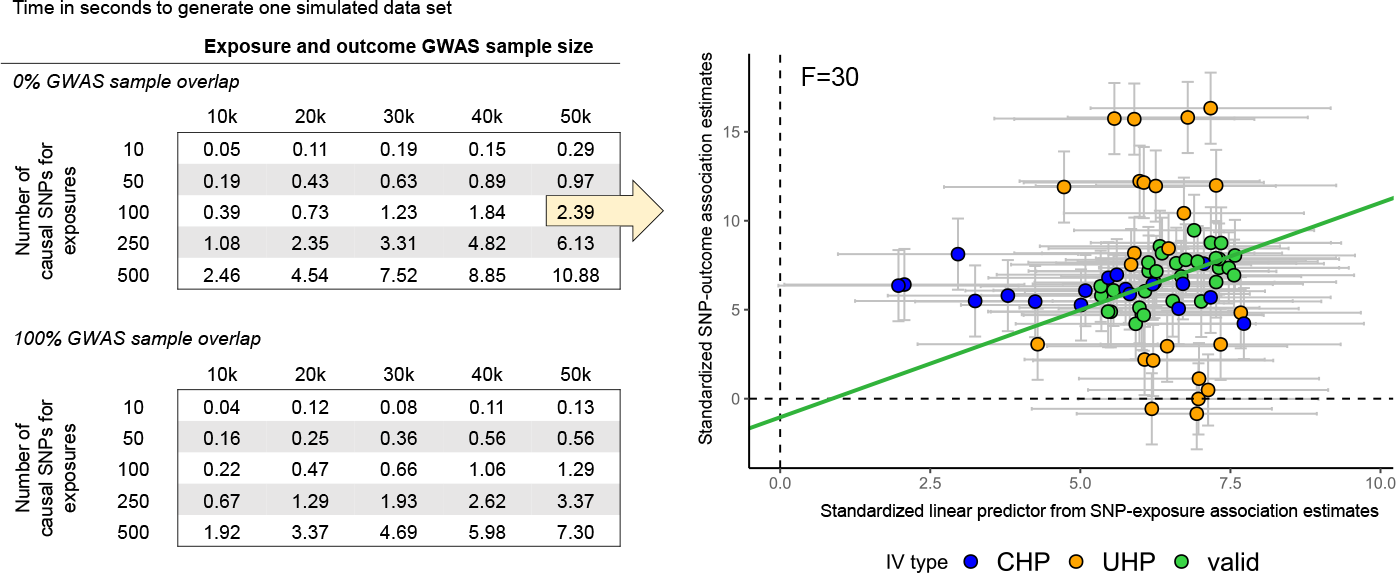
This figure shows computation times to generate one simulated data set using `simmr` (left) and an example of the generated summary data for evaluation of MR methods (right). The data in the right panel is only of the final selected and pruned set of instruments and was creating simply by executing `plot_simdata()` in R. The original number of causal SNPs was 100, but the selected number of IVs to achieve an F-statistic of 30 was 91. The full code used to produce this figure is available at https://github.com/noahlorinczcomi/simmr. The x-axis in the right panel is the standardized linear predictor used in multivariable MR 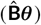 and the y-axis is the estimated association of the IVs with the outcome in standardized scale. The green line is the linear fit to the green points representing valid instrumental variables. The yellow arrow in the left panel indicates that the time it took to generate the data in the right panel was 2.4 seconds when there is no GWAS sample overlap and GWAS sample sizes are 50k. When the exposure and outcome GWAS sample overlap is 100%, computation time was reduced to 1.39 seconds.

### Algorithm 1

Pseudo-code of simulated data generation using `simmr`

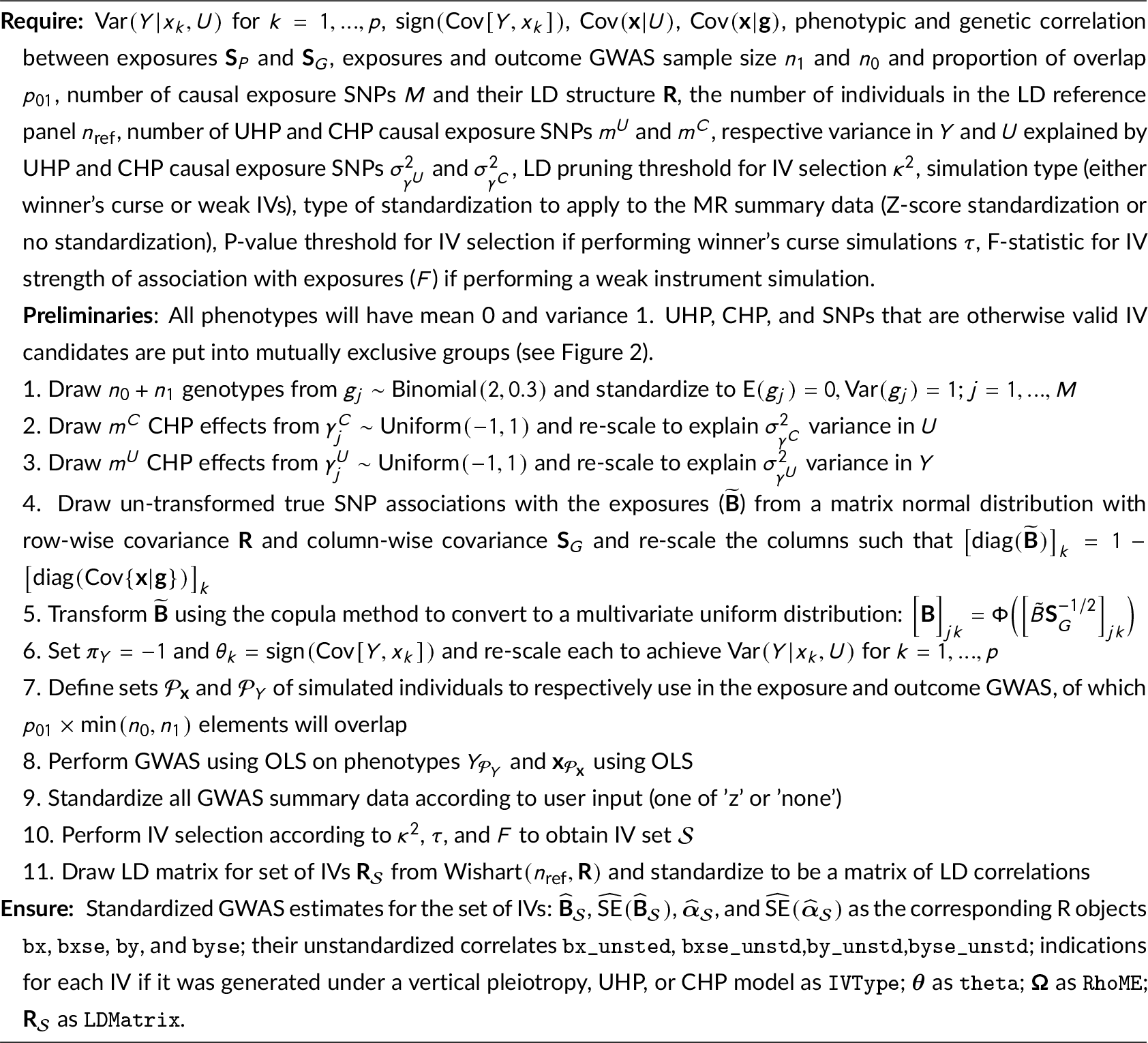

## 3 SOFTWARE

`simmr` uses the popular R software v4.1.1 or later (R Core Team, 2021) and only requires downloading 3 files: `basicfunctions.R`, `set_params.R`, and `generate_data.R`. Users of `simmr` change the parameters in the `set_params.R` file to their desired quantities. These parameters are displayed in blue text in Figure 2. Next, another file named `generate_data.R` is sourced and the simulation data is automatically loaded into the user’s R environment where simulations can be performed. The `simmr` software is available at https://github.com/noahlorinczcomi/simmr, where a tutorial is present and researchers can directly change any of the source code to improve it. Changes are tracked automatically and earlier versions can be restored at any time.

## 4 INPUT AND OUTPUT

Users of `simmr` input all parameters that are present in the `set_params.R` file. The output of executing the `generating_data.R` file after all parameters from `set_params.R` are present in the global environment is the following summary data: (i) standardized SNP association estimates between all IVs and each exposure in the R matrix object `bx`, (ii) their corresponding standardized standard errors in `bxse`, (iii) standardized estimates of SNP association between each IV and the outcome in the vector `by`, (iv) their corresponding standardized standard errors in `byse`, (v) all corresponding unstandardized estimates of SNP association between the exposures and outcome and their corresponding standard errors (`bx_unstd,bxse_unstd,by_unstd,byse_unstd`), (vi) the number of selected IVs in `mIVs`,an indication for each IV if it was generated under a non-UHP/CHP, UHP, or CHP model in the vector `IVType`,the matrix of estimated LD correlations between all IVs in the matrix `LD`, (ix) the true causal effects of each exposure on the outcome in the vector `theta`, (x) and the matrix of correlations between GWAS estimation errors for the outcome and all exposures in the matrix `RhoME`. Users can view a summary visualization of their simulated data by executing `plot_simdata()` after `generate_data()`.

## 5 CONCLUSION

There is currently no standard simulation framework for performing simulations in Mendelian Randomization research. Different researchers have independently performed simulations designed to reflect similar real world conditions, but the performance of the same methods can vary greatly. The MR literature is replete with MR estimators, with each at some point having a demonstrated advantage over others in simulation. The transferability of their performance to real world settings may be in question if the advantages can only be reached in specific conditions. We present `simmr`, an open-source software for performing simulations to evaluate the performance of Mendelian Randomization methods. Researchers can directly modify the `simmr`source code. It is our intention that the community will use this opportunity to establish an accepted procedure for performing simulations using MR. As `simmr` is refined and expanded, we expect it will provide a useful tool to facilitate future MR method development and evaluation.

## Abbreviations

MR: Mendelian Randomization;
LD: linkage disequilibrium;
UVMR: univariable Mendelian Randomization;
MVMR: multivariable Mendelian Randomization;
SNP: single nucleotide polymorphism;
GWAS: genome-wide association study;
DAG: directed acyclic graph

## Acknowledgements

This work was supported by grant HG011052 and HG011052-03S1 (to X.Z.) from the National Human Genome Research Institute (NHGRI)

## 6 APPENDIX

We mentioned in the Introduction section that at least 45 statistical methods for performing Mendelian Randomization (MR) exist. These methods include IVW (Burgess et al., 2013), dIVW (Ye et al., 2021), pIVW (Xu et al., 2022), MR-Egger (Bowden et al., 2016), MR-RAPS (Zhao et al., 2020b), MRAID (Yuan et al., 2022), MRMix (Qi and Chatterjee, 2019), MR-cML (Xue et al., 2021), MVMR-cML (Lin et al., 2023), MR-PRESSO (Verbanck et al., 2018), IMRP (Zhu et al., 2021), MR-Median (Bowden et al., 2016), MR-MaxLike (Burgess et al., 2016b), MR-Corr (Cheng et al., 2022a), MR-Robust (Rees et al., 2019), MR-Lasso (Kang et al., 2016), MR-Conmix (Burgess et al., 2020), CAUSE (Morrison et al., 2020), MR-CUE (Cheng et al., 2022b), MR-Horse (Grant and Burgess, 2023), MR-BMA (Zuber et al., 2020), MR-Robin (Gleason et al., 2020), EMIC (Jiang et al., 2022a), MR-Mode (Hartwig et al., 2017), MRBEE (Lorincz-Comi et al., 2023), MR-Lap (Mounier and Kutalik, 2023), the Wald test (Palmer et al., 2008), JAM (Newcombe et al., 2016), MR using factor analysis (Patel et al., 2023), mixIE (Lin et al., 2021), MRMO (Deng et al., 2022), BMRMO (Deng et al., 2023), BWMR (Zhao et al., 2020a), moPMR-Egger (Liu et al., 2021), sisVIVE (Kang et al., 2016), MR-LDP (Cheng et al., 2020), MR-CIP (Xu et al., 2021), MR-PATH (Iong et al., 2020), MR-Clust (Foley et al., 2021), BayesMR (Bucur et al., 2020), BMRE (Schmidt and Dudbridge, 2018), MR-link (van Der Graaf et al., 2020), OMR (Wang et al., 2021b), CoJo (Yang et al., 2012), MR using PCA (Burgess et al., 2017), and GRAPPLE (Wang et al., 2021a).

## Availability and requirements

**Project name**: `simmr`**Project home page**: https://github.com/noahlorinczcomi/simmr

**Operating system(s)**: Platform independent

**Programming language**: R

**Other requirements**: R 4.1.1 or higher; `mvnfast` (required) and `ggplot2` (optional) R packages

**License**: MIT

